# Nitrogen utilization and transformation of *Stenotrophomonas maltophilia* W-6 with nitrogen-fixing ability

**DOI:** 10.1101/337386

**Authors:** Shutong Wang, Yi Xu, Zhenlun Li

## Abstract

Strain W-6 was isolated from the purple soil and successfully identifed as *Stenotrophomonas maltophilia* and used for the investigation on nitrogen utilization. Strain W-6 was monitored with the ability of biological nitrogen fixation when N_2_ was used for the sole nitrogen source, and yet nitrogenase activity would be inhibited in the presence of extra nitrogen. Moreover, Strain W-6 could utilize NO_3_^−^, NO_2_^−^ and NH_4_^+^ for cell growth through assimilation, but unable to convert them to atmospheric nitrogen. Meantime, accumulation of nitrite was observed during the nitrate removal process, and the optimal conditions for nitrate removal were temperature of 20°C, shaking speed of 150 rpm, sodium succinate as the carbon source and C/N of 12. The experimental results indicate that *Stenotrophomonas maltophilia*utilize W-6 could utilize not only N_2_ but also other nitrogen sources directly as its N substance. Therefore, heterotrophic *Azotobacter* may possess a great significance to nitrogen cycle except in biological nitrogen fixation.

**Importance:** *Azotobacter* spp. are found in soils worldwide, with features not simply for the nitrogen fixation, but for the energy metabolism relevant to agriculture. However, the role of *Azotobacter* potential in the function of nitrogen cycle except in biological nitrogen fixation is largely unknown. As such, whether bacteria utilize either inorganic nitrogen or organic nitrogen has remained obscure. The present studies indicate that *Stenotrophomonas maltophilia* W-6 could highly efficient utilize nitrate, nitrite and ammonium etc. N substance and detect NH_4_^+^ as final product. The transport velocities of nitrate-N to nitrite-N was quickly without gaseous nitrogen was produced. We probed the relationship between biological nitrogen fixation and N cycle via N conversion processes by *S. maltophilia* W-6 with nitrogen-fixing ability

## Introduction

Nitrogen is one of the most abundant and pervading element on the face of the earth, but most organisms, plants and animals included, are unable to access atmospheric dinitrogen for metabolic purposes (1). **B**iological **n**itrogen **f**ixation (BNF) by means of prokaryotic metabolic process responsible for converting the most abundant but relatively inert form of N into biologically available substrates is the main way for introduction of N into the biosphere. Therefore, the higher nitrogenase activity in the organisms especially *Azotobacter* play an important role in nitrogen recycles and is a primary N input pathway in many ecosystems and sustains global plant productivity (2).

Conventional biological fixation of nitrogen conversion atmospheric nitrogen to ammonium was believed to be inhibited that takes place after supplying nitrogen fertilizer to the soil (3). Besides, high concentration of NH_4_^+^ have bad effect on the growth of microbe and their normal physic-chemical functions (4). Nevertheless, the removal and utilization of nitrogen by bacteria are affected significantly by many factors. However, due to the complexity of the soil environment, some correlative mechanism and effect factors need to be further researched. To date, no one has isolated pure bacteria to study the nitrogen transformation mechanism of *Azotobacter* under different nitrogen conditions and conducted in-depth studies on nitrogen cycling pathways of *Azotobacter.* So far, the species of *A.chroococcum* are the most prevalent species found in *Azotobacter* but other species described including *A.agilis* (5), *A.vinelandii* (6), *A.beijerinckii* (7), *A.insignis* (8), *A.macrocytogenes* (9) and *A.paspali* (10) according to the reports published. Though, as gram negative bacteria, *Stenotrophomonas maltophilia* has received a lot of attention in the current virus researches (11,12), its capacity as *Azotobacter* for the utilization and conversion of different nitrogen has not been studied a lot.

In the present study, several studies were under way to determine whether external nitrogen sources (NO_3_^−^, NO_2_^−^, NH_4_^+^, NH_2_OH and organic nitrogen) have effect on *Azotobacter*-related law of utilization and transformation of nitrogen. In addition, whether the concentration of the nitrogen will affect the properties of the *Azotobacter* hasn’t been reported yet. In this investigation, we obtained a strain *Stenotrophomonas maltophilia* which have relatively few studies been done of its use as *Azotobacter.* We investigated the utilization and transformation of different nitrogen in Stenotrophomonas maltophilia W-6 with nitrogen-fixing ability and further characterized the key factors which affect N utilization efficiency under aerobic conditions. Therefore, the research was essential to further reveal great significance of *Azotobacter* not only about biological nitrogen fixation but about the nitrogen cycle.

## Results

### Screening and Identification of the Studied Bacterium

By repeated streaking on fresh agar plates, the bacterium with aerobic heterotrophicnitrogen fixation capability was obtained. The strain W-6 could growth on the Ashby nitrogen-free medium, accompanied with TN increased. It was successfully identified based on the analysis of 16S rRNA gene sequence analysis as *Stenotrophomonas maltophilia* and the phylogenetic tree was successfully constructed (**Fig. 1**). Besides, through the detection of the nitrogenase gene, it is successfully depicted that the strain W-6 belongs to *Azotobacter*. (**Fig.2**). Therefore, strain W-6 could be identified as *Azotobacter* based on the TN increased and the analysis of 16s rRNA and nitrogenase gene.

**Fig 1.**
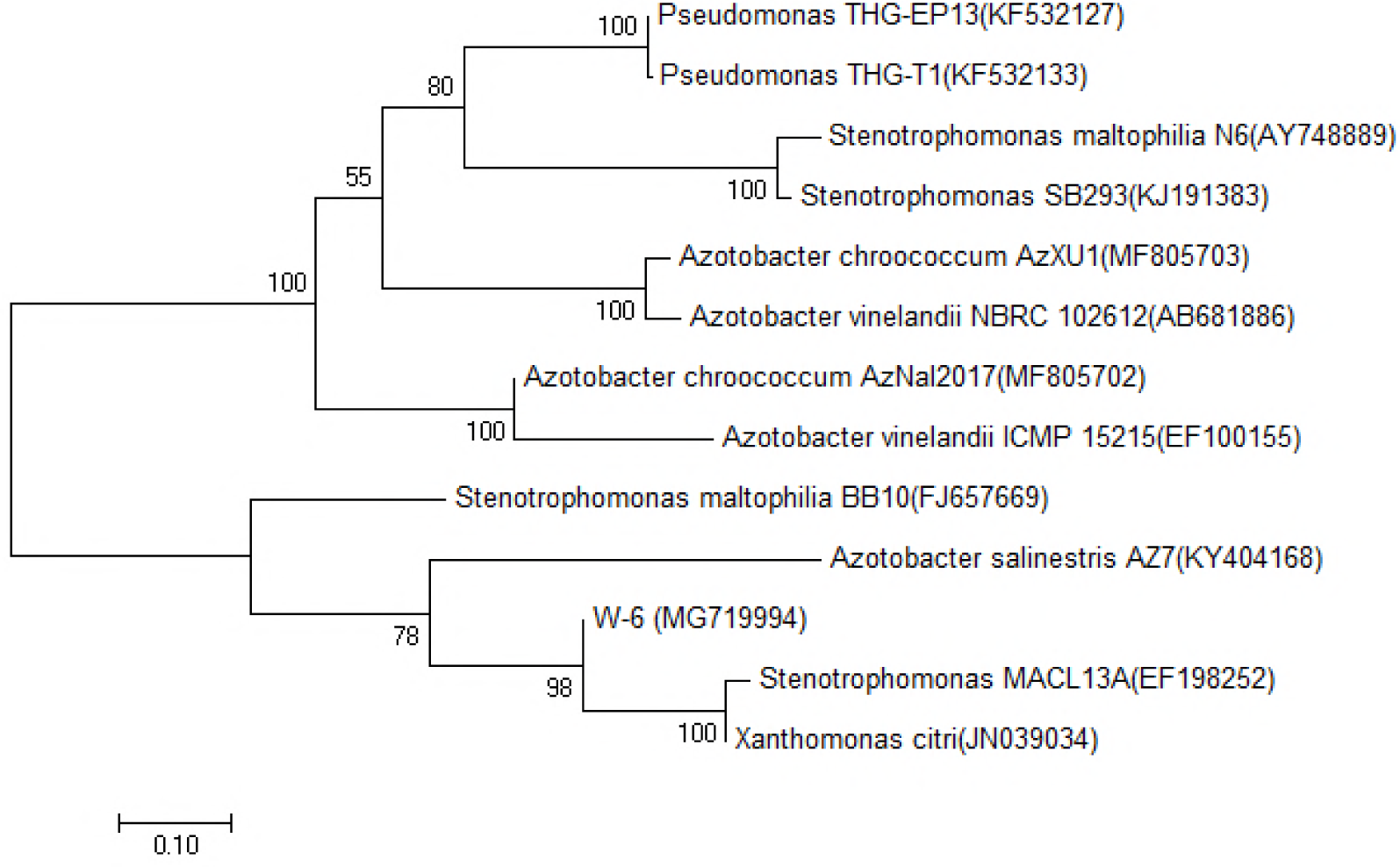
The phylogenetic tree of W-6 based on 16S rRNA gene sequence analysis.

### Exploration of utilization and transformation of nitrate of W-6 under low concentration of nitrate nitrogen

The nitrate utilization and transformation activity of W-6 was obviously shown the change in TN, NO_3_^−^-N and OD_600_ with low concentration of nitrate nitrogen (7.05 mg/L) supplied as the nitrogen source (**Fig. 3**). While initial N utilization quickly, nitrogenase activity was inhibited. After the external nitrogen source was used up, W-6 started to fix nitrogen itself which shown a significant nitrogen-fixing capacity.

**Fig 2.**
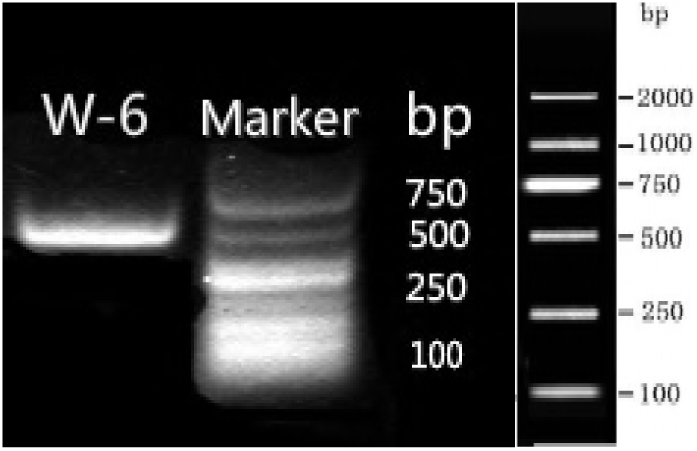
The PCR amplification of Nitrogenase gene in W-6. (M-2000 bp DNA ladder)

**Fig 3.**
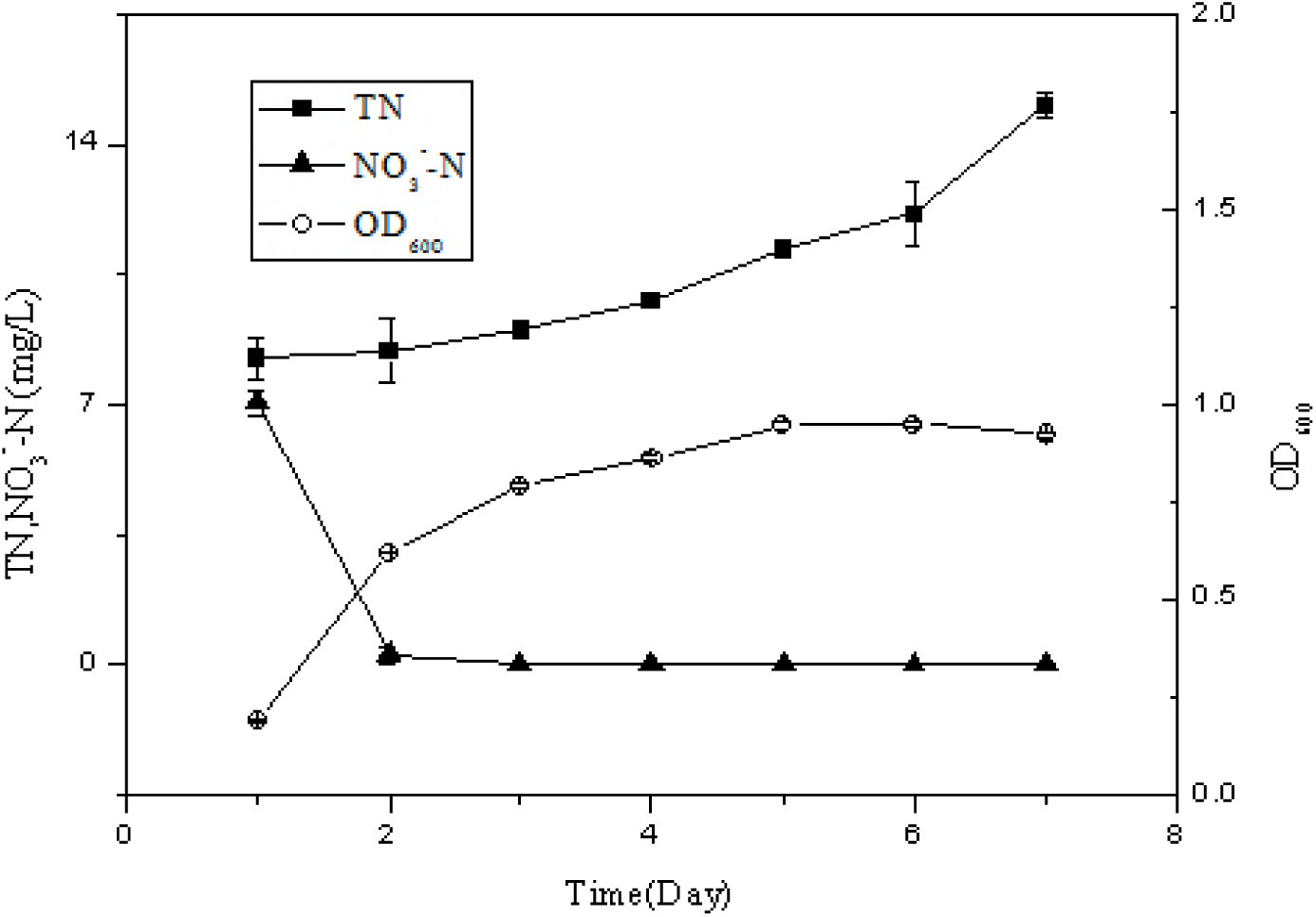
Growth activity and TN accumulation performance of strain W-6 in the low concentration of nitrate nitrogen media.

### Ammonium Removal Performance of Strain W-6

Several survey results indicate that ammonium have a great significance for the exploration of nitrogen cycling. Therefore, to further elucidate the ammonium nitrogen conversion process in water was necessary to evaluate effect of the conversion of ammonium on strain W-6. The result for the test has shown that NH_4_^+^-N was completely removed in 18 h by strain W-6 using 148.81 mg/L initial concentration (**Fig. 4**). Meanwhile, NO_3_^−^-N and NO_2_^−^-N were undetected during the experiments means that strain W-6 could not invert ammonium to nitrite to nitrate (NH_4_^+^ → NO_3_^−^ → NO_2_^−^).

**Fig 4.**
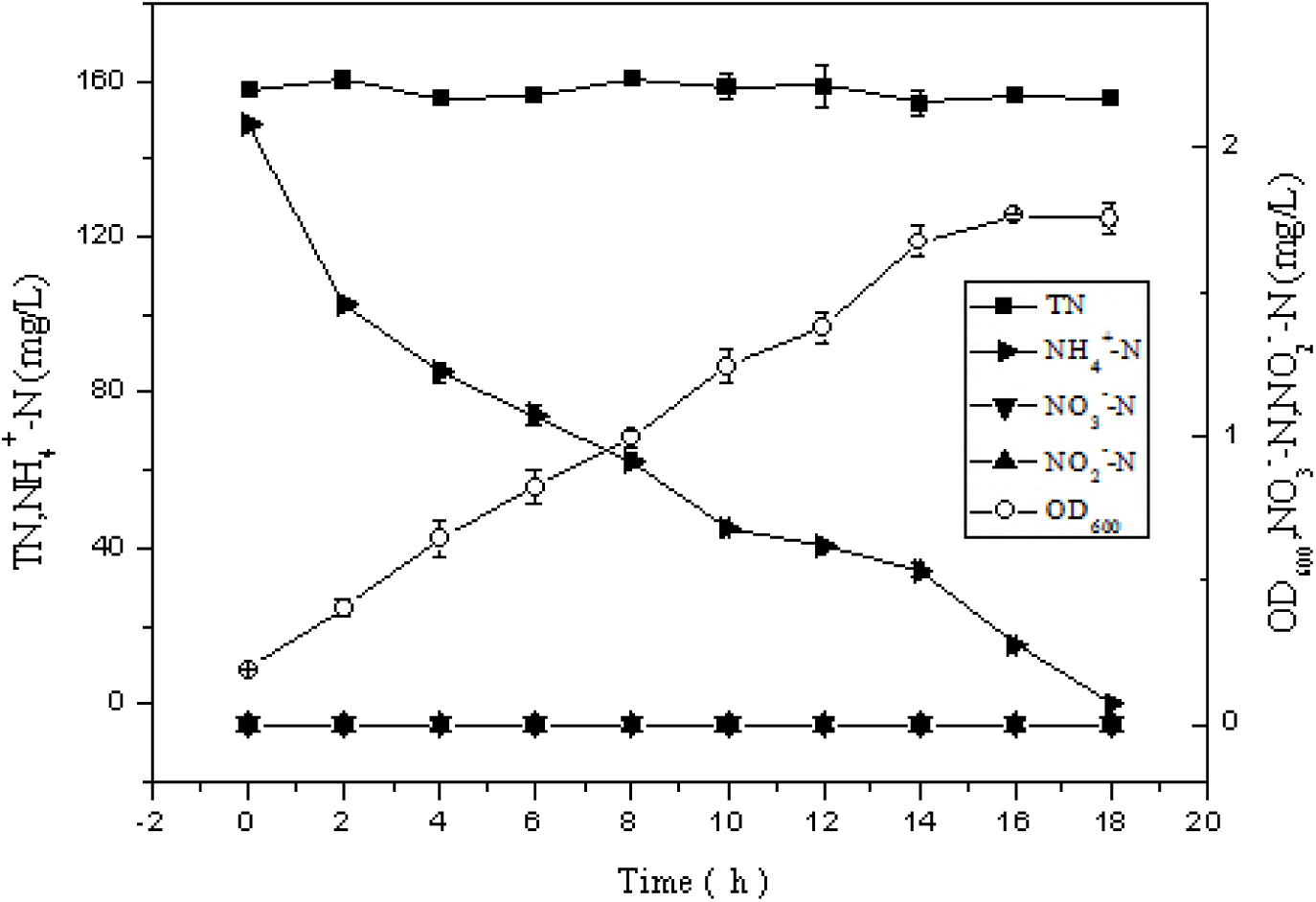
Growth activity and ammonium conversion performance of strain W-6 in the ammonium nitrogen media.

### Nitrate Removal Performance of Strain W-6

The line chart was made showed fate of utilization and conversion by strain W-6 with nitrate supplied as the sole nitrogen source (**Fig. 5**). It merits our attention that NO_3_^−^-N was almost completely removed in 55 h using 100.90 mg/L initial nitrate concentration. Simultaneously, NH_4_^−^-N as nitrogen conversion product was detected increasing in the whole process especially during stationary phase. The results would indicate that the final conversion that nitrate nitrogen to ammonium nitrogen is one of the pathways for ammonium assimilation in W-6.

**Fig 5.**
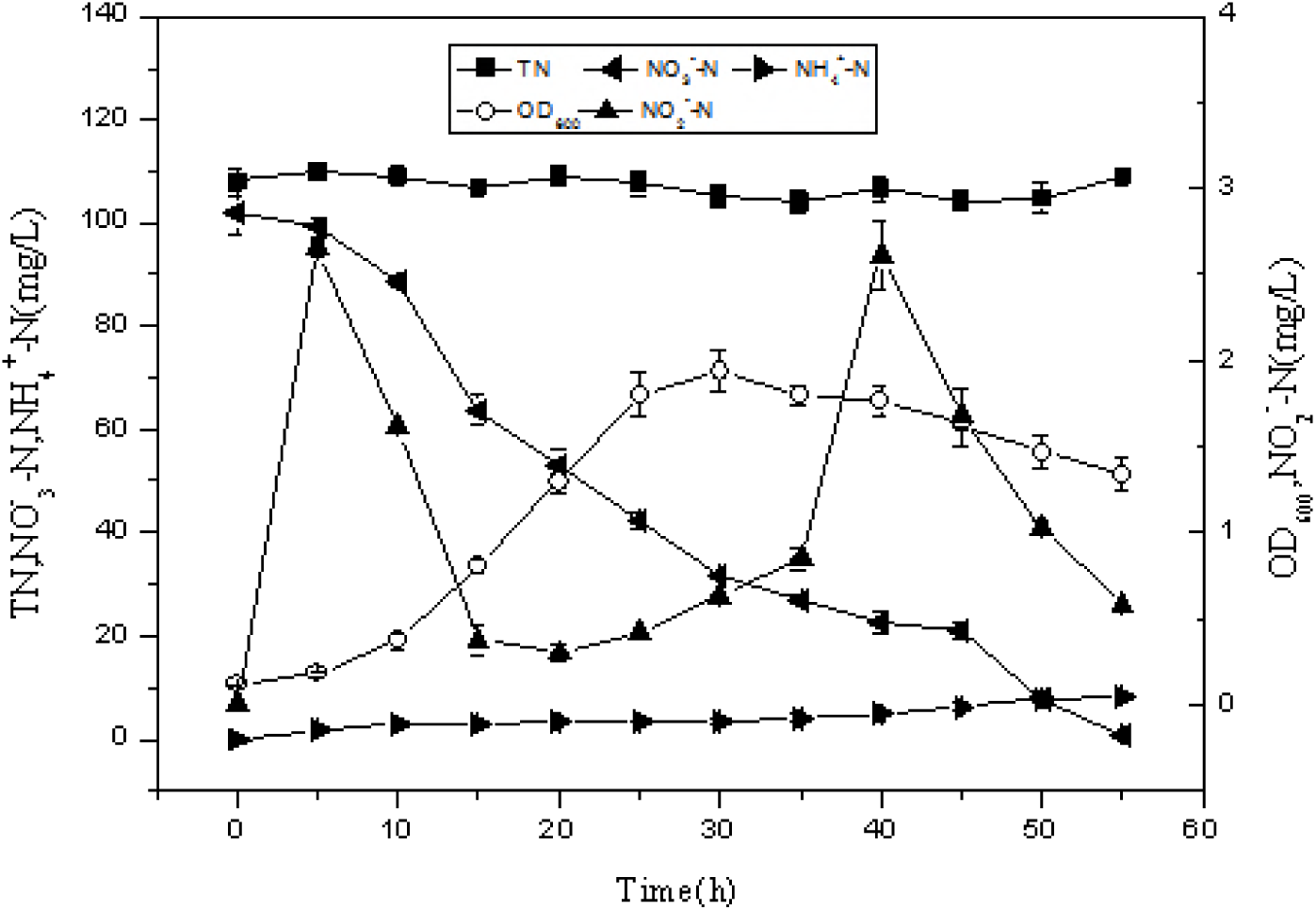
Characteristics of N conversion and utilize by strain W-6 with nitrate nitrogen supplied as the sole nitrogen source.

Beside ammonium was produced after the decomposition of organic nitrogen into ammonium in the body and then lysed after the death of the microorganism. Meanwhile, the other product nitrite nitrogen experienced two fluctuations, one increased after the initial increase, another increased latter and then decreased to substantially none. This might reveal the transformation pathway of N element in microbial metabolism

### Nitrite Removal Performance of Strain W-6

Nitrite is an important intermediate, no matter in the process of nitrification or nitrification. The results depicted the conversion of nitrogen that nitrite was transformed and utilized 83.17 % within 60 h (**Fig. 6**). It’s worth noting that nitrate nitrogen was not detected during the entire process, indicating that W-6 does not have the ability to convert nitrite nitrogen to nitrate nitrogen. In addition, ammonium nitrogen was detected 17.93 mg/L in 60 h. Moreover, ammonium production was almost absent at the beginning of growth. After W-6 growth rapidly in the fourth day, the production of ammonium also increased significantly. It could explain that strain W-6 produces ammonium due to cells autolytic lysis after death once again.

**Fig 6.**
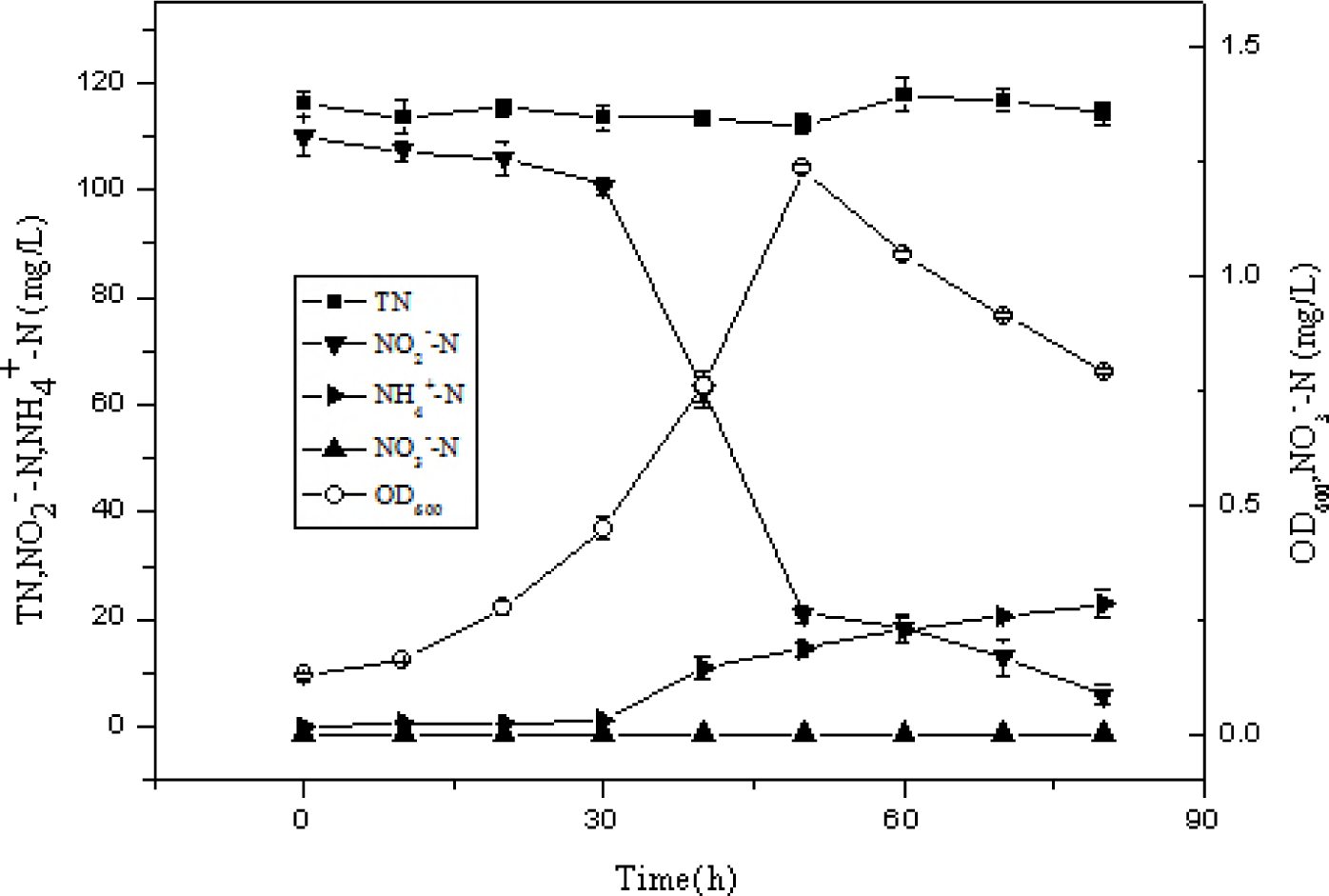
Characteristics of N conversion and utilize by strain W-6 with nitrite nitrogen supplied as the sole nitrogen source.

### Assessment of Organic Nitrogen Removal Performance by Strain W-6

N in the soil existed mainly in the form of organic nitrogen, therefore, to investigate the effects of organic nitrogen on *Azotobacter* makes sense. The results indicated variation trend of organic nitrogen utilized and ammonium by strain W-6 with organic nitrogen (**Fig. 7**). Strain W-6 could make good use of organic nitrogen and ammonium was produced simultaneously. Of particular concern is ammonium was also produced in large quantities in anaphase of the state period or prophase of the death period. It has further proven the conclusion mentioned before.

**Fig 7.**
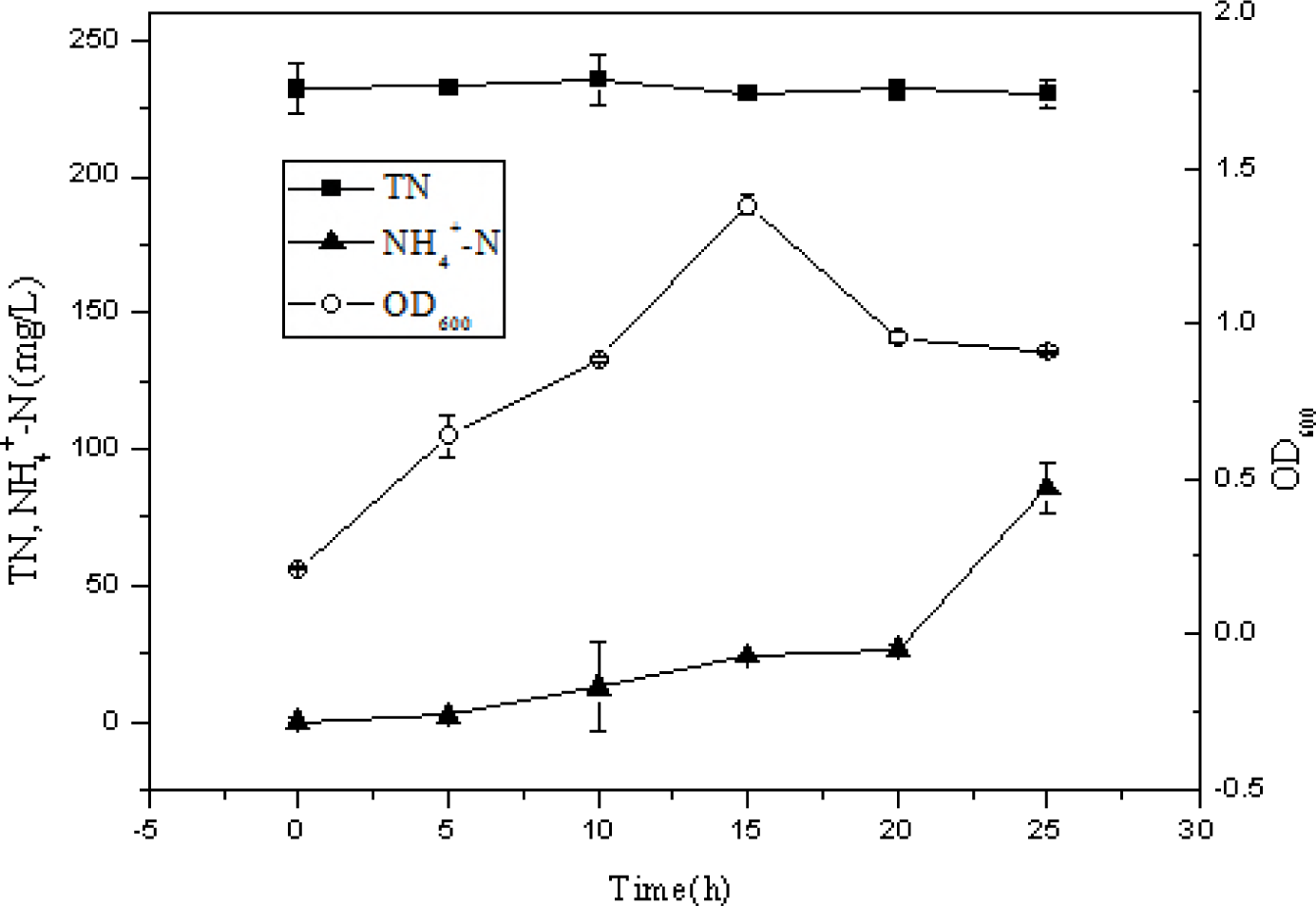
Characteristics of N conversion and utilize by strain W-6 with nitrite nitrogen supplied as the sole nitrogen source.

### Assessment of Simultaneous Growth Curve and Hydroxylamine Utilization and Conversion Performance of Strain W-6

To assess the growth curve and conversion and utilize capability of strain W-6 with hydroxylamine was used as a single nitrogen source. The results were obviously shown that strain W-6 grew rarely slowly even its viability level was restrained at the first 3 d and OD_600_ reached 0.15 from 0.18 mg/L (**Fig. 8**). After 8 d of incubation, the concentration of hydroxylamine nitrogen decreased infinitesimal from 15.36 mg/L to 9.54 mg/L, and only 37.87 % of hydroxylamine removal was achieved. Furthermore, it was attractive that nitrate nitrogen was detected little and nitrite nitrogen was detected infinitesimal. This would be an interest found that HAO might exist in W-6 which has the function of oxidizing hydroxylamine to nitrite.

**Fig 8.**
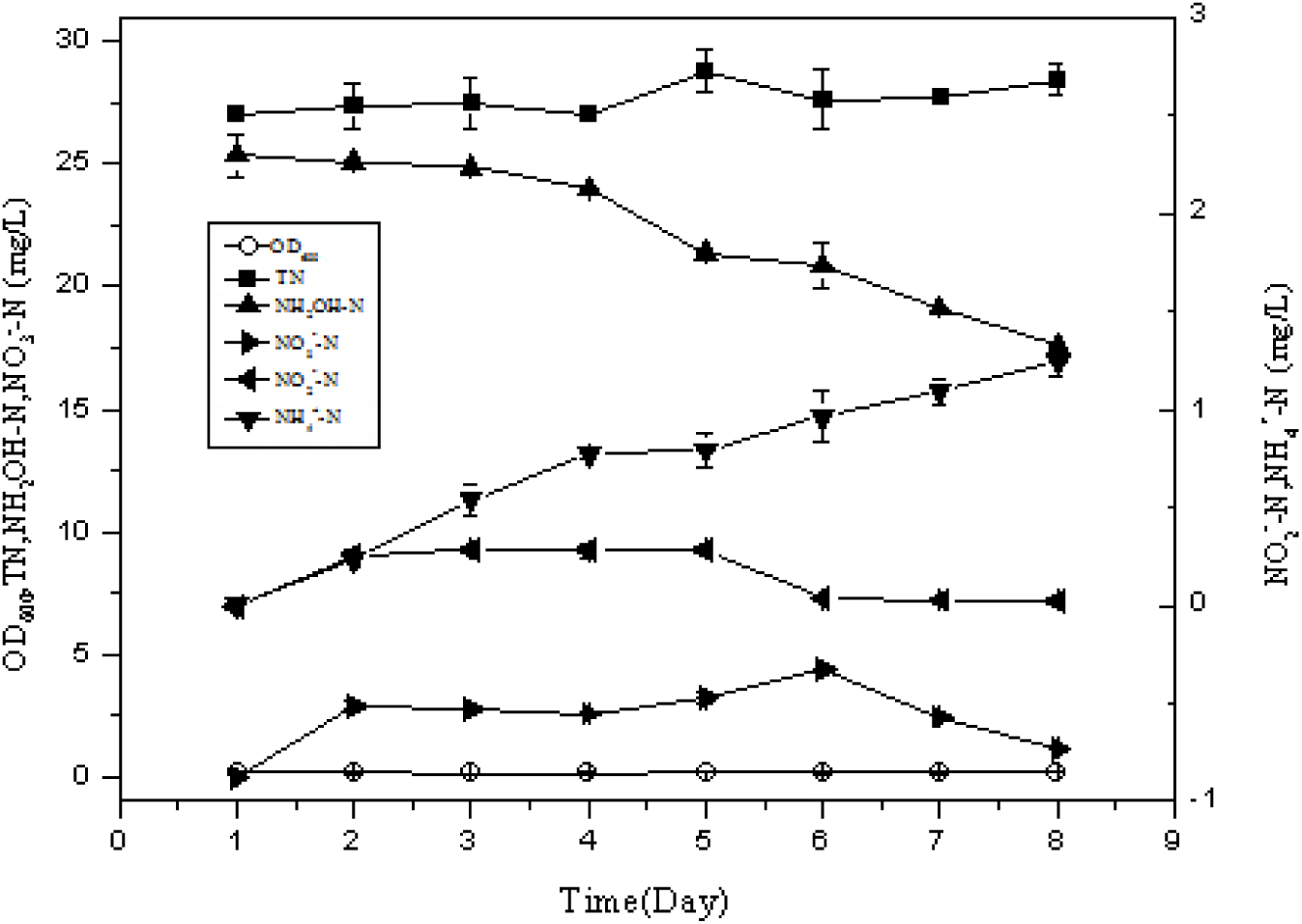
Characteristics of N conversion and utilize by strain W-6 with hydroxylamine *supplied as the sole nitrogen source*.

### Assessment of Different Concentration of Ammonium Removal Performance

It is generally known that high concentration of ammonium is toxic to the microbial. Strain W-6 showed growth activity with different high concentration of ammonium nitrogen (about 300, 600, 900, 1200, 1500 and 1800 mg/L) supplied as nitrogen sources (**Fig. 9**). It is visible that W-6 peaked at 2.78 mg/L in 5 d when the concentration of ammonium was 300 mg/L. In other situations, it was screened growth continuously for the initial 6 d, while the highest value was reached at 5.28 mg/L when ammonium concentration was 600 mg/L. Meantime, W-6 was able to tolerate as high as 1200 mg/L ammonium.

**Fig 9.**
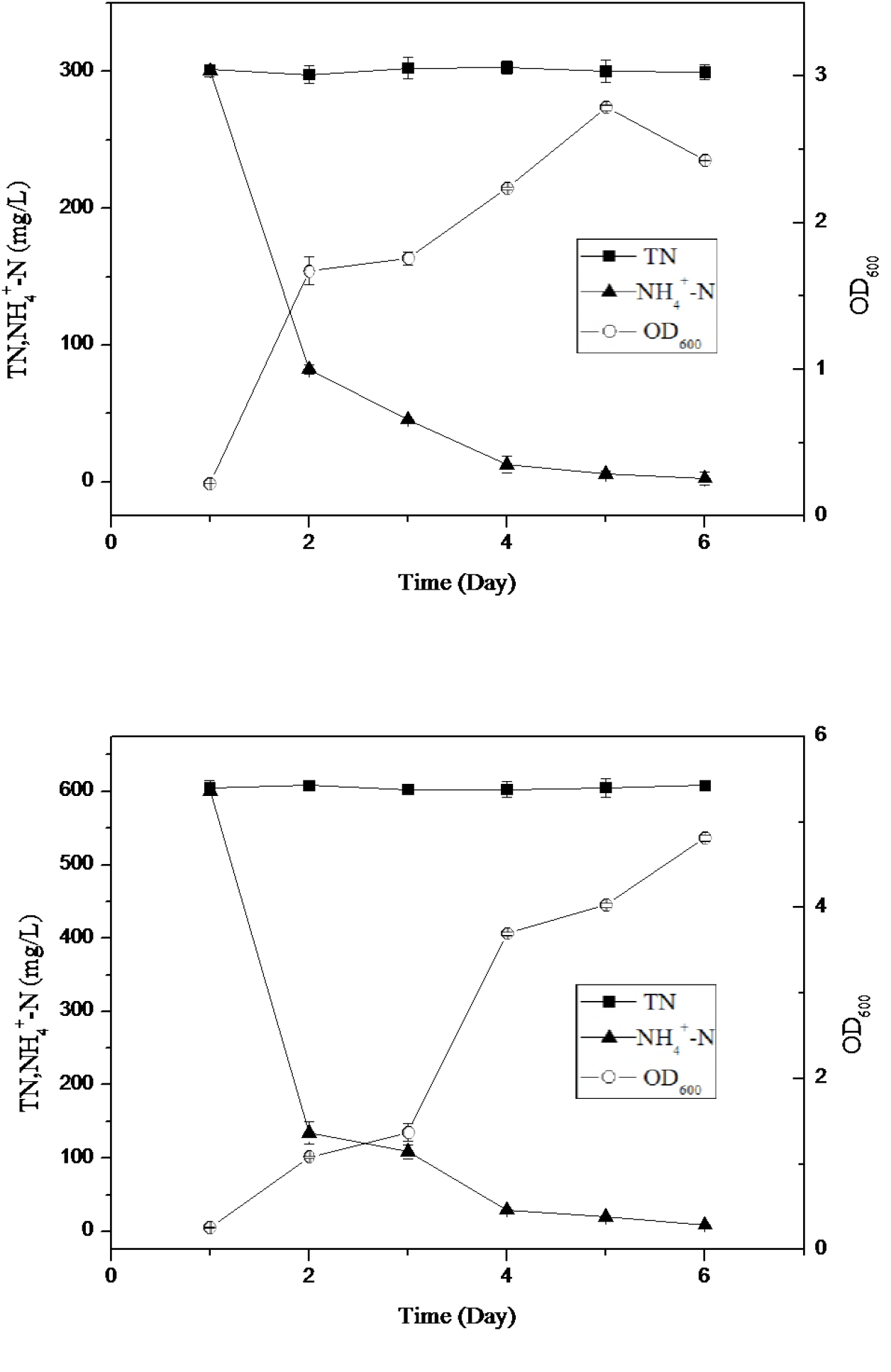
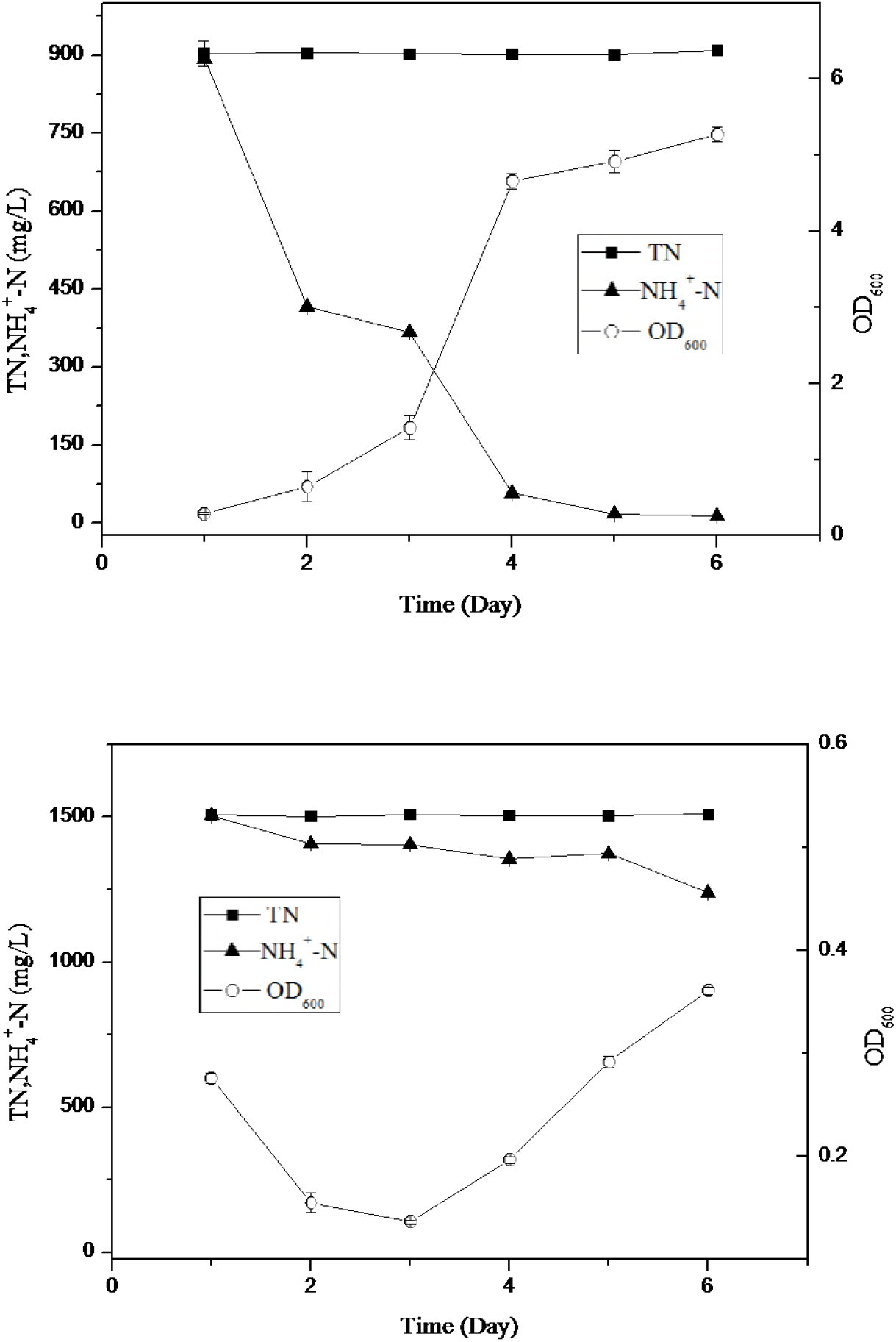
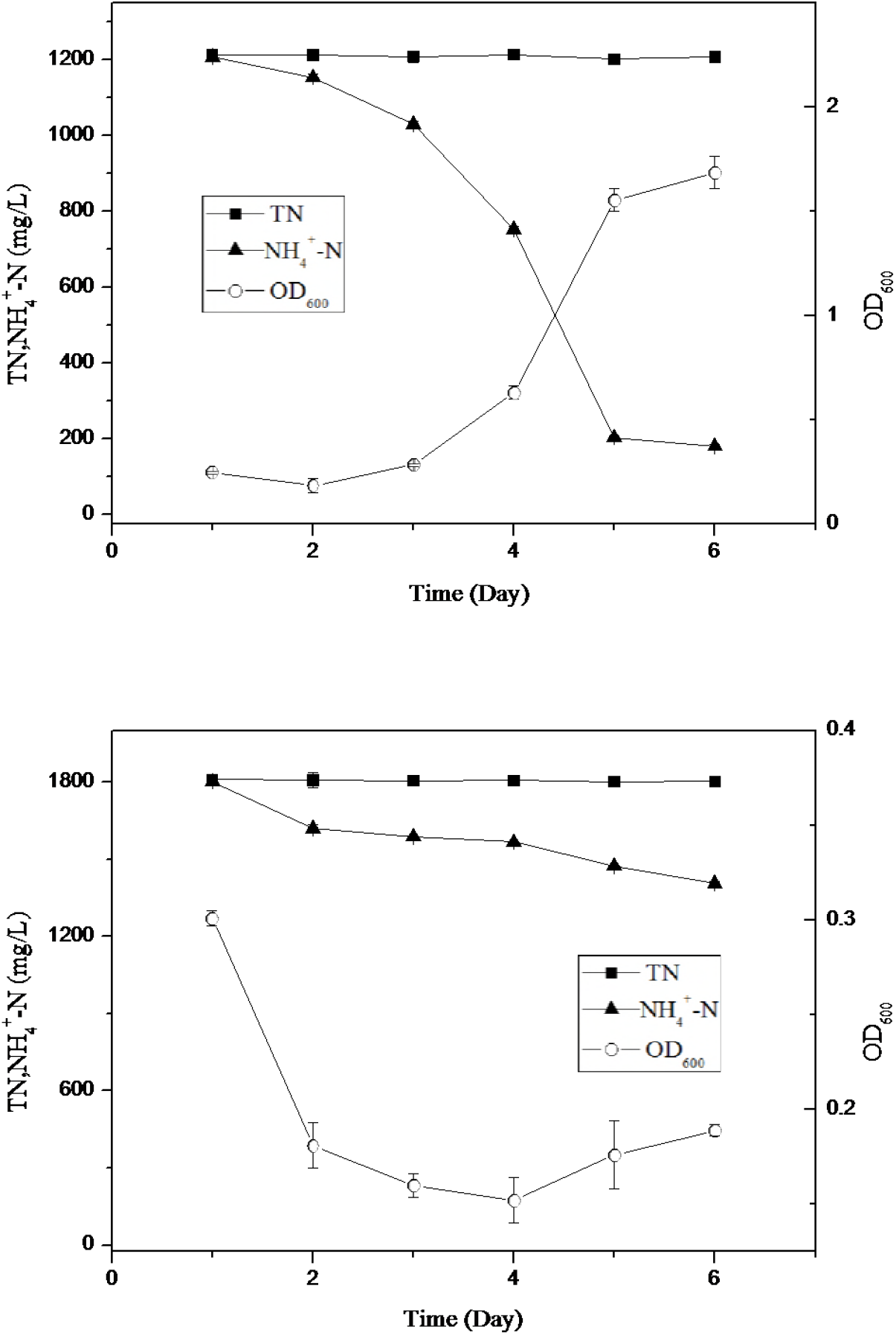
Characteristics of N conversion and utilize by strain W-6 with different high concentrations of ammonium nitrogen supplied as the nitrogen source.

Additionally, it depicts nitrogen removal characteristics of strain W-6 at different high concentrations of ammonium nitrogen under aerobic conditions at 20°C. When strain W-6 was cultivated at the initial ammonium nitrogen concentrations of 300.52, 600.38, 893.02,1206.85, 1504.86 and 1801.93mg/L for 6 d, the corresponding ammonium removal efficiencies were 99.13 %, 98.50 %, 98.49 %,84.98 %, and 24.24 % and 21.97 %, respectively.

### Exploration of the Influence of Different Carbon Source on Nitrate Utilization

The growth activity, especially nitrate removal characteristic of strain W-6 was used as important indicators to determine the optimal carbon source. Compared with other carbon sources, a significant decrease of nitrate was observed from sodium succinate as the OD_600_ increased from 0.13 to 1.92 mg/L, and removed approximately 99.64 % of nitrate nitrogen (61.564 mg/L initial NO_3_^−^-N) at the same time (**Table 1**).

**Table 1:**
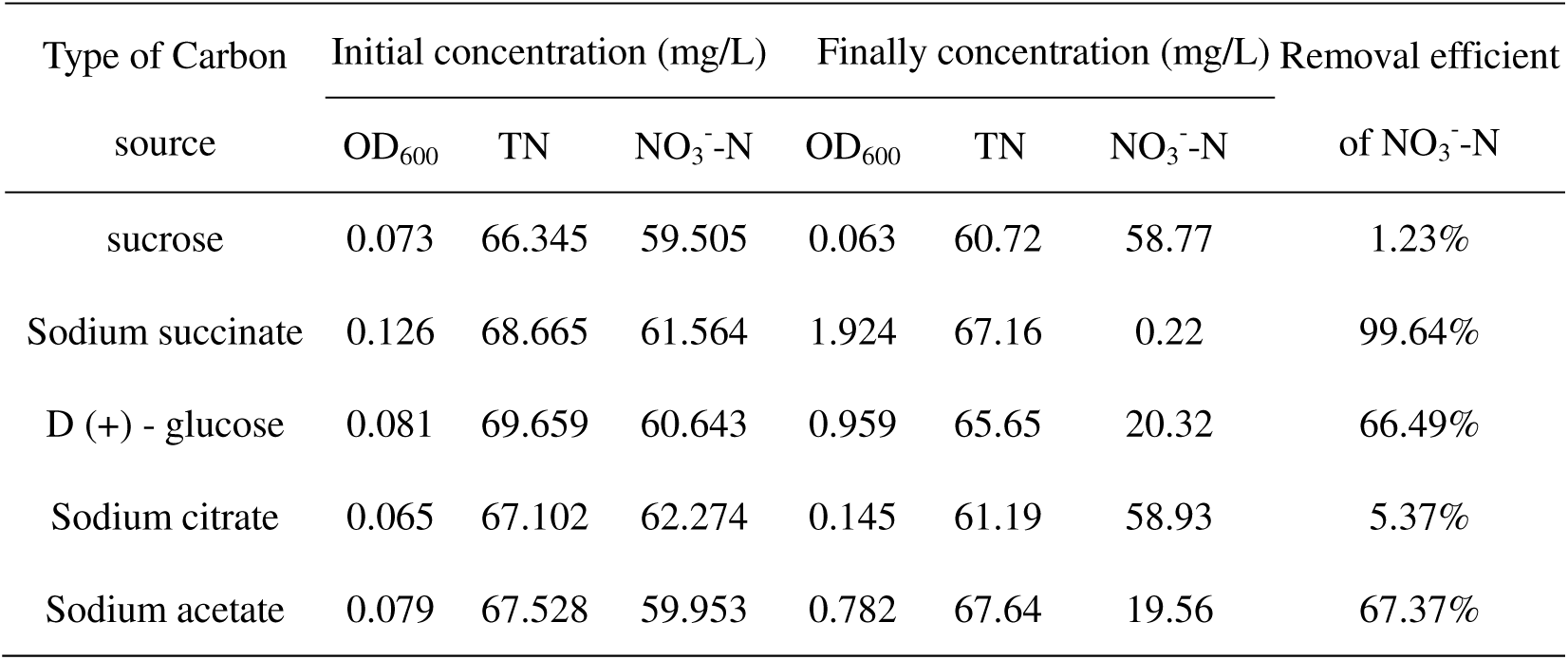
*Growth activity and NO*_*3*_^*-*^*-N removal performance of strain W-6*

### Exploration of the Influence of Different Temperature on Nitrate Utilization

Generally, the suitable temperature for the growth of microorganisms is about 30°C. The maximum occurring at 34-35°C and the suggested positions of the upper and lower limits of respiration at about 50 and 10°C. A recently reported nitrogen-fixing bacterium with high nitrate reduction and ammonium assimilation has an optimum culture condition of 25°C (13). In contrast, 20°C resulted in maximum nitrate removal efficiency among the temperatures tested by W-6. In the context of that the initial inoculum volume is basically the same as the nitrate nitrogen concentration, the bacterial growth is faster and the nitrate-nitrogen use efficiency is highest at 20°C, reached 82.24 % (**Table 2**).

**Table 2:** *Growth activity and NO*_*3*_^*-*^*-N removal performance of strain W-6*

| Type of<br>temperature | Initial concentration (mg/L) | | | Final concentration (mg/L) | | | Removal efficient<br>of $\text{NO}_3^-$-N |
| --- | --- | --- | --- | --- | --- | --- | --- |
| | $\text{OD}_{600}$ | $\text{NO}_3^-$-N | TN | $\text{OD}_{600}$ | $\text{NO}_3^-$-N | TN | |
| 15°C | 0.083 | 54.75 | 61.19 | 0.561 | 23.57 | 60.06 | 56.95% |
| 20°C | 0.080 | 54.53 | 61.34 | 0.973 | 9.68 | 60.55 | 82.24% |
| 25°C | 0.075 | 54.48 | 60.91 | 0.923 | 10.85 | 60.77 | 80.09% |
| 30°C | 0.089 | 54.16 | 61.57 | 0.848 | 14.79 | 61.19 | 72.69% |

### Exploration of the Influence of Different Carbon and Nitrogen Ratio on Nitrate Utilization

Both carbon and nitrogen are inaccessible factors in the growth and development of life and play important roles in the structural composition and metabolism of the organism. By experiment the C/N ratio to explore the influence of these factors on the nitrate utilization. The effect of different level of carbon nitrogen ratios by strain W-6 is investigated. Among the carbon nitrogen ratios tested, the highest bacterial growth activity and nitrate nitrogen removal rate was achieved when the carbon to nitrogen ratio is 12. The results showed that C/N is 12 was better than others for the efficient of NO_3_^−^-N removal efficiency reached 98.58 % (**Table 3**).

**Table 3:** *Growth activity and NO*_*3*_^*-*^*-N removal performance of strain W-6*

| Type of C/N | Initial concentration (mg/L) | | | Final concentration (mg/L) | | | Removal efficient<br>of $\text{NO}_3^-$-N |
| --- | --- | --- | --- | --- | --- | --- | --- |
| | $\text{OD}_{600}$ | $\text{NO}_3^-$-N | TN | $\text{OD}_{600}$ | $\text{NO}_3^-$-N | TN | |
| 0 | 0.186 | 58.46 | 77.81 | 0.172 | 55.01 | 78.43 | 5.91% |
| 3 | 0.179 | 59.74 | 71.14 | 0.347 | 42.93 | 70.33 | 28.13% |
| 6 | 0.185 | 61.83 | 71.23 | 0.534 | 26.80 | 73.41 | 56.65% |
| 9 | 0.188 | 61.13 | 74.97 | 1.101 | 5.04 | 74.17 | 91.75% |
| 12 | 0.179 | 61.0 | 74.83 | 1.413 | 0.86 | 71.89 | 98.58% |
| 15 | 0.189 | 60.78 | 73.69 | 0.872 | 1.88 | 73.17 | 96.90% |
| 18 | 0.197 | 60.20 | 66.88 | 0.718 | 14.67 | 68.63 | 75.63% |

### Exploration of the Influence of Different Dissolved Oxygen on Nitrate Utilization

The amount of oxygen is important for the metabolism of microorganisms. In the laboratory, we simulated the environmental conditions suitable for the growth of bacteria, and used a constant temperature shaker to maintain a suitable oxygen input, so that the microbial growth environment can reach the optimum conditions. The speed for growth activity and NO_3_^−^-N removal performance by W-6 was conducted. The results showed that the activity of growth and nitrate nitrogen removal along with different speeds. The maximum activity was obtained when speed is 150 rpm. It is obviously showed that W-6 reached 0.47 mg/L from 0.05 mg/L and removed 75.38 % of nitrate (**Table 4**).

**Table 4:** *Growth activity and NO*_*3*_^*-*^*-N removal performance of strain W-6*

| Rotating<br>speed/rpm | Initial concentration (mg/L) | | | Final concentration (mg/L) | | | Removal efficient<br>of $\text{NO}_3$-N |
| --- | --- | --- | --- | --- | --- | --- | --- |
| | $\text{OD}_{600}$ | $\text{NO}_3$-N | TN | $\text{OD}_{600}$ | $\text{NO}_3$-N | TN | |
| 0 | 0.041 | 56.92 | 60.19 | 0.125 | 19.31 | 60.17 | 66.07% |
| 50 | 0.054 | 54.83 | 61.09 | 0.226 | 19.28 | 60.71 | 64.83% |
| 100 | 0.080 | 55.14 | 61.75 | 0.487 | 19.75 | 59.67 | 64.19% |
| 150 | 0.053 | 54.94 | 61.33 | 0.467 | 13.52 | 61.47 | 75.38% |
| 200 | 0.052 | 55.06 | 61.47 | 0.477 | 16.03 | 60.71 | 70.88% |
| 250 | 0.088 | 57.96 | 61.47 | 0.417 | 23.06 | 61.04 | 60.22% |

## Discussion

Generally speaking, over high concentration of ammonium (beyond 100 mg/L) is toxic to organisms (4). Up to now, only few bacteria including *Acinetobacter junii* YB (14), *Alcaligenes faecalis* No.4 (15) and *Vibrio* sp. Y1-5 (16) exhibited capable of adapting to high ammonium concentration and performing high ammonium resistance characteristics were isolated. In this research, strain W-6 showed strong resistance to ammonium at a concentration of 1200 mg/L. The results demonstrated that *Azotobacter* W-6 could utilize high concentrations of ammonium for metabolism. In addition, it is noted that this research is the first to prove the *Stenotrophomonas maltophilia* capable of growing well on medium containing high concentrations of ammonium (more than 1200 mg/L). When strain W-6 was cultivated at the initial ammonium nitrogen concentrations of 300.52, 600.36, 893.02, 1206.85, 1504.86 and 1801.93 mg/L for 6 d, the corresponding ammonium removal efficiency were 99.13 %, 98.50 %, 98.49 %, 84.98 %, and 24.24 % and 21.97 % respectively. That is, high concentrations of ammonium could be utilized and transformed by strain W-6 and used.

The previous study proved that when cells grown in the presence of some of inorganic nitrogen (NO_3_^−^, NO_2_^−^ and NH_4_^+^), nitrogenase activity was undetectable (17). Conversely, nitrogenase activity developed after the cells were cultured in the nitrogen-free medium. Nitrogen-fixing ability of *Azotobacter* was inhibited when external nitrogen sources were added had been reported (18), indicated that the microorganism gave priority to the use of an additional nitrogen source even at low concentrations. In a surprise, *Azotobacter* W-6 could utilize addition of a low concentration of nitrogen source for metabolism. Different results had been reported on *Azotobacter chroococcm* ATCC-4412 (18) in which couldn’t perform normal nitrogen fixation when using 0.22 mM KNO3. The reasons for formation this phenomenon possibility due to the fact that strain W-6 could use nitrogen quickly, and immediately fix N after nitrogen source was used up. The intermediates undetected during the nitrogen removal process in *Azotobacter* W-6 require further investigation.

Ammonium (NH_4_^+^), nitrite (NO_2_^−^) and nitrate NO_3_^−^) are the most common ionic forms of dissolved inorganic nitrogen in aquatic ecosystems (19). In this research, strain W-6 could utilized inorganic nitrogen (NO_3_^−^, NO_2_^−^ and NH_4_^+^)and organic N at 20°C. The removal efficiencies of nitrate and nitrite were 92.31 % and 80.85 % in 50 h when the initial concentrations were 101.76 and 109.89 mg/L. Meanwhile, 148.81mg/L ammonium nitrogen was completely reduced in 18 h. Figure 2 showed that nitrite and nitrate were undetected during ammonium oxidation and, most likely, W-6 doesn’t have denitrification capacity. Similar result had been reported on *Pseudomonas tolaasii* Y-11(20) in which nitrite and nitrate were not observed during ammonium oxidation, either.

It merits our attention that hydroxylamine couldn’t be directly utilized by *Stenotrophomonas maltophilia* W-6 for assimilation unless converted into nitrite. Results from figure 8 indicated that nitrite was detected when hydroxylamine was used as nitrogen source, which had similar result with Otte (21). Apparently, NH_2_OH induced HAO activity in strain W-6. So the result explained that though without nitrification, strain W-6 might also have HAO.

Ammonium could be detected as final product with any kind of nitrogen source described the same result by Willis (22). Experimental results showed that only trace amounts of ammonium were produced in the initial growth period, the most were present in anaphase of the state period and prophase of the death period. This phenomenon showed that W-6 absorbed inorganic nitrogen converted into organic nitrogen in the body and released ammonium after the bacteria have died. However, different results have been reported, such as the process of dissimilatory nitrite reduction to ammonium occurred in strain Y-9 (23).

There is also an interesting phenomenon worthy of our attention is that the pH of the medium we finally measured were about 8.5 from the initial about 7.0 after every performed task. This may be due to the release of OH^−^ in the metabolic process of *Azotobacter,* which leads to a rise in pH. It’s was reported in *Thioalkalivibrio versutus*, strain AL2 (22). Whereas *A.vinelandii* was able to produce acid which was different from W-6 (24). Further research is needed before we can draw firm conclusions.

As a result, the phylogenetic tree of W-6 was constructed based on 16S rRNA gene sequence analysis as shown in figure 1 (GenBank Accession Number MG719994). The result obviously showed that strain W-6 was clustered with *Stenotrophomonas, Xanthomonas citri* while diverged from *A.chroococcum, A.vinelandii* and *Pseudomonas*. The tree suggests that W-6 and *Stenotrophomonas, Xanthomonas citri* may have relatively close phylogenetic relationship comparing to *A.chroococcum, A.vinelandii* and *Pseudomonas*. It merits our attention that though W-6 has nitrogen fixation capacity, it phylogenetic relationship is very far from the typical *Azotobacter* such as *A.chroococcum and A.vinelandii.* It is likely that the W-6 has undergone a gene mutation during the evolution of the system, and the *Stenotrophomonas* mutation generates a corresponding gene that can fix nitrogen. However, what kind of gene or which locus has been mutated and what kind of mutation has occurred requires further study.

In this research, *Stenotrophomonas maltophilia* W-6 was able to utilize many kinds of inorganic (NO_3_^−^, NO_2_^−^, NH_4_^+^ and NH_2_OH) and could tolerate high concentration of ammonium (over 1200 mg/L). The absorption capability to ammonia of W-6 is the strongest much more than the lowest hydroxylamine. Meanwhile, the accumulation of ammonium mainly appeared in both present in anaphase of the state period or prophase of the death period. This research is the first comprehensive research closely reported the use of different nitrogen from the perspective of pure bacteria about *Azotobacter* and may possess a great significance to nitrogen cycle. The further investigation of W-6 should be on the mechanisms of nitrogenase to learn more about the nitrogen fixation characteristics of *Azotobacter*. By genetically analyzing the origin of the strain, it is also worth investigating the source of its variation.

## Materials and Methods

### Media

Ashby Nitrogen-free medium(25): Mannitol 1 %, KH_2_PO_4_ 0.02 %, MgSO4·7H _2_O 0.02 %, NaCl 0.02 %, CaSO_4_·2H _2_O 0.01 %, CaCO_3_ 0.5 %. Solid medium should plus agar 8 %. Ashby Nitrogen-free medium was used to isolate *Azotobacter*.

Lysogeny broth (LB) medium (26)(per liter): 5 g of yeast extract, 1 g of peptone, 5 g of NaCl, pH 7.0-7.2 (added 2 % of agar if it is solid).

The basal medium used for bacteria cultivation contained (per liter):0.1 g of MgSO4·7H _2_O, 7.9 g of K_2_HPO_4_·3H _2_O, 1.5 g of KH_2_PO4 and 2 ml of trace elements solution. The kinds and concentrations of nitrogen sources were adjusted for different test requirements.

Trace element solution(27)(per liter): 1.5 g of FeCl_3_·6H _2_O, 0.15 g of H_3_BO_3_, 0.03 g of CuSO_4_·5H _2_O, 0.03 g of KI, 0.06 g of Na_2_MoO_4_·2H _2_O, 0.12 g of MnCl_2_·4H _2_O, 0.12 g of ZnSO4·7H2O, 0.12 g of CoCl _2_·2H _2_O.

All biochemical reagents except LB were of analytical grade.

### Isolation of bacteria

In order to the isolation of bacteria, 5 g of soil sampling was transferred to sterile Ashby nitrogen-free culture medium(25) in 250 ml flasks and agitated on the temperature controlled rotary shaker to gain homogeneous suspensions (20°C, 150rpm, 24h). Then the suspended liquid (5ml) was pipetted into Ashby Nitrogen-free medium for *Azotobacter* accumulation and incubated under the conditions described above for secondary isolate. In order to gain pure *Azotobacter*, this one step needs to be repeated at least 5 times. Finally, single and purified colony was obtained by isolating and repeatedly plating 3 times on agar containing Ashby Nitrogen-free culture medium named W-6. The strain W-6 was suspended in 25 % glycerol solution at −80 °C for long-term storage(28).

### Identification by 16S rRNA gene sequence analysis

16S rRNA technology has been considered a very mature technology in the identification of species(29). Primers were used to amplify 16S rRNA gene by a PCR protocol (PTC200, Bio-Rad, USA). Partial 16S rRNA genes were sequenced by Shanghai Invitrogen Biotech Company Ltd (Shanghai, China). Finally, phylogenetic tree for 16S rRNA analysis was constructed based on the partial 16S rRNA sequence of strainW-6. All sequences were aligned using MEGA 7.0 program and a phylogenetic tree was built in MEGA 7.0 using the neighbor-joining method(30). Moreover, nitrogenase-stabilized gene fragments were also detected by cloning-PCR. These primers were 35F (5′-AAA GGY GGW ATC GGY AAR TCC AC-3′) and 491R (5′-TTG TTS GCS GCR TAC ATS GCC ATC AT-3′) (6,31).

### Assessment of nitrogen removal capability with different nitrogen source

In order to determine the utilization and transformation of different nitrogen in strain W-6, single colony was cultivated for 24 h in 100 ml sterile Lysogeny broth medium(26)at 20°C, 150 rpm. Then the strain cell in initial cultivation medium was harvested by centrifuging at 4000 rpm for 8 min and the precipitate washed twice with sterile water. Then, the pellets were inoculated into 100 ml media which were added nitrate, nitrite, ammonium, hydroxylamine, organic nitrogen as nitrogen sources in BM medium respectively. In addition, the pellets were inoculated into 100 ml media which were added different concentration of ammonium in BM medium. They all use sodium succinate as carbon source in the condition that the carbon to nitrogen ratio is 12. Besides, the medium without inoculation was used as control. All experiments were carried out in triplicate.

### Effects of external factors on the removal efficiency of nitrate experiments

External factors such as carbon source, C/N ratio, temperature and dissolved oxygen seem to influence the removal efficiency of nitrate. Over the course of the study, strain W-6 was incubated in Lysogeny broth medium at 20°C with a shaking speed of 150 rpm until the OD_600_ reached approximately 1.0 and the aggregates of microbe were obtained by centrifugation (6000 rpm, 8 min). Then the pellets were inoculated into 100 ml BM medium using nitrate as nitrogen source. Likewise, the medium without inoculation was used as control and experiments were carried out in triplicate.

### Analytical measurements and data analysis

Most indicators were measured using a spectrophotometer (DU800, Beckman Coulter, USA) except for pH. The cell density was monitored as OD_600_ was measured at a wavelength of 600nm. Total nitrogen (TN) concentration in suspension was analyzed using alkaline potassium persulfate digestion and its corresponding absorbance value was calculated by the equation: A=A_220_-2A_275_ to eliminate or reduce the impact of background factors. Nitrate, nitrite and ammonium were determined by standard methods after centrifugation. The concentrations of NH_4_^+^-N, NO_3_^−^-N and NO_2_^−^-N were analyzed using the supernatant with different wavelength. NH_4_^+^-N concentration was determined using indophenol blue colorimetry with a wavelength 625nm. NO_3_^−^-N concentration associated absorbance determination was the similar as TN and calculated using the same method. NO_2_^−^-N concentration was determined at a wavelength of 540 nm after adding 1 mL ofchromogenicreagentin20min.NH_2_Ohwasmeasured by indirect spectrophotometry(32).

Statistical analysis and graphic plotting were conducted by using SPSS Statistic, Excel, and Origin 8.6. For each kind of medium, the results are presented as means ± SD (standard deviation of means).

## Acknowledgments

This work was supported financially by the National Key Research and Developmental Program of China (2017YFC0404705)

